# Computational prediction of the effect of amino acid changes on the binding affinity between SARS-CoV-2 spike protein and the human ACE2 receptor

**DOI:** 10.1101/2021.03.24.436885

**Authors:** Chen Chen, Veda Sheersh Boorla, Deepro Banerjee, Ratul Chowdhury, Victoria S Cavener, Ruth H Nissly, Abhinay Gontu, Nina R Boyle, Kurt Vandergrift, Meera Surendran Nair, Suresh V Kuchipudi, Costas D. Maranas

## Abstract

The association of the receptor binding domain (RBD) of SARS-CoV-2 viral spike with human angiotensin converting enzyme (hACE2) represents the first required step for viral entry. Amino acid changes in the RBD have been implicated with increased infectivity and potential for immune evasion. Reliably predicting the effect of amino acid changes in the ability of the RBD to interact more strongly with the hACE2 receptor can help assess the public health implications and the potential for spillover and adaptation into other animals. Here, we introduce a two-step framework that first relies on 48 independent 4-ns molecular dynamics (MD) trajectories of RBD-hACE2 variants to collect binding energy terms decomposed into Coulombic, covalent, van der Waals, lipophilic, generalized Born electrostatic solvation, hydrogen-bonding, π-π packing and self-contact correction terms. The second step implements a neural network to classify and quantitatively predict binding affinity using the decomposed energy terms as descriptors. The computational base achieves an accuracy of 82.2% in terms of correctly classifying single amino-acid substitution variants of the RBD as worsening or improving binding affinity for hACE2 and a correlation coefficient r of 0.69 between predicted and experimentally calculated binding affinities. Both metrics are calculated using a 5-fold cross validation test. Our method thus sets up a framework for effectively screening binding affinity change with unknown single and multiple amino-acid changes. This can be a very valuable tool to predict host adaptation and zoonotic spillover of current and future SARS-CoV-2 variants.

## Introduction

The ongoing coronavirus disease 2019 (COVID-19) pandemic caused by the severe acute respiratory syndrome coronavirus 2 (SARS-CoV-2) continues to be a major global challenge to public health and has caused unprecedented losses to the global economy^1^ and ecology^2^. Multiple vaccines have received emergency use authorization (Pfizer, Moderna and J&J) and additional vaccines are under Phase-3 clinical trials (AstraZeneca, Janssen and Novavax) in the United States. However, several new variants of the wild-type (WT) virus (i.e., isolate Wuhan-Hu-1, GenBank: MN908947) have emerged in United Kingdom^4^ (B.1.1.7 lineage), South Africa^5^ (B.1.351 lineage), and Brazil^6^ (P.1 lineage) with increasing prevalence worldwide.^7^ A few non-synonymous mutations leading to amino-acid changes in the spike protein of these variants are associated with enhanced infectivity. The enhanced infectivity of the variants is possibly through increased binding affinity of the receptor binding domain (RBD) of the spike protein with the human angiotensin converting enzyme-2 (hACE2) receptor^8^ and through changes in the conformational dynamics of the spike protein.^9^ Although a recent preliminary report suggests that the current vaccines can still effectively protect people from SARS-CoV-2 variants^10^, another report showed that plasma from recipients of Moderna (mRNA-1273) or Pfizer-BioNTech (BNT162b2) vaccines was significantly less effective in neutralizing SARS-CoV-2 variants encoding E484K or N501Y or the K417N:E484K:N501Y combination.^11^ In addition, a significant decrease in neutralizing titers against the B.1.351 but not the B.1.1.7 UK variant, with plasma from mRNA-1273 vaccinated humans and non-human primates has been observed.^12^ Hence, continued surveillance and methods to accurately predict affinity gains of the RBD-hACE2 binding event due to amino acid changes in the RBD are extremely important.

SARS-CoV-2 is an enveloped virus with a single-stranded RNA genome of approximately 30 kb size^13^. The mutation rates of RNA viruses upon replication are generally higher than DNA viruses, which could be as high as 10^−4^ to 10^−3^ per nucleotide incorporated ^14^ SARS-CoV-2 has a mutation rate of on average 7.23 mutations per sample^15^ which is significantly lower than HIV and Influenza-A viruses.^16,17^ The simultaneous incorporation of multiple (i.e., 15-20) amino-acid changes in a few emerging strains such as ΔFVI (Danish mink), B.1.1.7 (UK), B.1.1.54 (SA) are a cause of concern as it suggests further adaptation of the virus and fitness gains in humans and other animals.^18,19,20^ Several of these variants involve amino acid changes in the spike protein suspected to increase transmissibility^21^, alter infectivity^22,23^ and/or escape neutralizing antibodies.^22,24^ The viral spike protein binding to the hACE2 protein is the first and crucial step in viral entry.^25–31^ The spike protein makes contact with hACE2 using 16 residues of the 223 amino acid long receptor binding domain (RBD) forming multiple polar and hydrophobic interactions.^32^ The binding strength between RBD and hACE2 thus directly affects infection dynamics and potentially disease progression. Starr et al.^33^ exhaustively assessed the impact of single amino acid changes in the RBD of the SARS-CoV-2 quantifying the effects on RBD expression and hACE2 binding. It was revealed that most amino acid changes (i.e., 84.5%) are detrimental for RBD expression and hACE2 binding, around 7.5% of mutations are neutral but about 8% enhance hACE2 binding. The corresponding amino acid changes in RBD that lead to enhancements in binding with hACE2 can potentially become additive in their contribution to receptor affinity. Even though the RBD accounts for only 2% of the amino acid changes observed in the entire spike protein^34^, it is the target for more than 90% of the neutralization antibodies generated by humoral response^35^. Therefore, RBD is the most susceptible target to antigenic escape by amino acid changes. Consequently, amino acid changes in RBD that can increase binding affinity with hACE2 and/or adversely affect antibody neutralization have been extensively mapped by high-throughput mutational studies.^36,37^ For example, the amino acid change Y453F in the RBD present in the ΔFVI (Danish mink) variant increases the binding affinity to hACE2 by four fold^38^ while also managing to partially evade the monoclonal antibody REGN10933 present in the Regeneron antibody cocktail.^39^ These studies highlight the importance of monitoring single and multiple amino acid changes in the spike RBD variants and their potential for increased binding affinity with ACE2 and/or immune escape. It is important to note that binding of the viral RBD with the hACE2 receptor is a necessary step but is not sufficient to cause a productive viral infection. Proteolytic cleavage of S1/S2 and S2’ sites is also needed to expose the fusion peptide enabling membrane fusion followed by viral entry at the surface or upon endocytosis.^40^ Furthermore, the host cellular environment must be permissive to viral RNA genome replication, translation to proteins and assembly into new virions.^41^

Computational methods can help assess the mechanistic role of the amino acid changes occurring in circulating viral variants and also predict potentially problematic amino acid changes that have not been identified so far. In a recent study, Chowdhury et al.^32^ biophysically characterized the binding interactions of human ACE2 with SARS-CoV-2 and SARS-CoV, uncovering the molecular details associated with the increased infectivity of CoV-2, relative to CoV. In another effort, Mohammad et al.^42^ calculated that the D614G variant has a higher computational binding interaction energy with furin. This was later experimentally corroborated revealing that the D614G change both increases RBD accessibility to binding with hACE2^43^ and enhances the efficiency of furin cleavage.^44^ Zhou et al.^24^ performed molecular dynamics (MD) simulations and molecular mechanics/Poisson-Boltzmann surface area (MM-PBSA) analysis on the N439K variant suggesting a higher binding affinity to hACE2 and resistance to the antibody REGN10987. These findings were supported by experimental evidence for the N439K variant escaping several neutralizing antibodies including REGN10987^24^. Several studies focus on testing the effect of one or several key single mutations, but systematic methods to predict and analyze a wider multi-mutational landscape are still lacking. It is worth noting that Chen^45^ et al. used an algebraic topology-based machine learning model to quantify the binding free energy changes of RBD from several existing CoV-2 variants. However, the performance of the method used was tested on the general SKEMPI-2.0^46^ dataset and not on a SARS-CoV-2 specific dataset.

Several computational approaches have been developed to predict the effect of amino-acid substitutions on protein-protein binding affinity. Some of them directly use energies from molecular-mechanics based empirical force-fields like FoldX^47^ and Rosetta^48,49^ or energies from molecular mechanics – generalized Born surface area (MM-GBSA) analysis of ensembles obtained from MD simulations^50^. Other methods such as SAAMBE^51^ and BindProfX^52^, use a combination of physical energies and residue-level structural properties or sequence-based conservation profiles, respectively. Purely statistical potentials like BeAtMuSiC^53^ and Contact potentials^54^ have also been explored. Updating the weights of energy terms using experimentally determined ΔΔ*G*_*bind*_ defined as the change in the free energy of binding upon amino-acid changes (i.e., ΔΔ*G*_*bind*_ = Δ*G*_*variant*_ − Δ*G*_*WT*_) has been shown to improve the prediction performance^55,56^ of molecular mechanics-based force-fields such as Rosetta^49^. The recently introduced machine learning (ML) based method TopNetTree achieves a better correlation coefficient over several existing methods on two benchmark datasets AB-Bind and SKEMPI.^57^ In spite of the existence of many different ΔΔ*G*_*bind*_ prediction methods, as reviewed recently^58^, performance is not always robust on unseen datasets not part of training data. The major limiting factor contributing to test set prediction inaccuracies is the paucity of experimental datasets (on ΔΔ*G*_*bind*_) with good coverage of both types and locations of amino acid changes.^58^

In this work, we first tested the predictive power of both parameterized force fields (i.e., Rosetta^49,59^) and detailed MM-GBSA^60–62^ analysis of MD simulation trajectories with explicit water molecules treatment ^63,64^ (see Table 1) in reproducing experimental RBD-hACE2 binding affinity data reported by Starr et al.^33^ Predictions using the MM-GBSA binding energies provided only partial agreement with experimental data (i.e., r = 0.33). Therefore, we next used experimental RBD-hACE2 binding energy terms to train a neural network (NN) regression model (NN_MM-GBSA) using the decomposed MM-GBSA energy terms as features and the experimental dissociation constants (*K*_D,app_) ratio between mutated variants and the wild-type as the regression target. Figure 3 pictorially illustrates the computational pipeline employed for building the model.

**Table 1.** Comparison of prediction performance of Rosetta and MM-GBSA with the regression models trained on MM-GBSA energies. *The standard deviation obtained for the five repetitions of the five-fold cross validation and training is shown within the parentheses.

| Method | Correlation coefficient (r) | % Variant correct recovery (VC) |
| --- | --- | --- |
| Rosetta | 0.47 | 68.52 |
| MM-GBSA | 0.33 | 61.11 |
| NN_ MM-GBSA* | 0.69 (0.03) | 82.24 (1.98) |
| Linear_regression_MM-GBSA* | 0.53 (0.17) | 59.43 (7.50) |
| NN_Rosetta* | 0.56 (0.08) | 74.03 (2.75) |

Agreement between experiment and the NN_MM-GBSA model predictions were significantly better than raw MM-GBSA energies reaching a correlation coefficient of r = 0.69 and an accuracy of 82.24% in recovering of the effect of amino-acid changes (i.e., in classifying as improving or worsening the binding affinity). The NN_MM-GBSA model also provided good predictions on binding affinities of the RBD from currently circulating SARS-CoV-2 variants (see Table 2). The achieved accuracy of prediction makes this model a useful tool for the computational assessment of current/emerging CoV-2 variants. The MM-GBSA energies and the NN_MM-GBSA model are available in Github at https://github.com/maranasgroup/NN_MM-GBSA_CoV2.

**Table 2.** Predictions of *K*_D,app_ ratios for single amino-acid changes found in circulating strains of SARS-CoV-2 and for two novel variants predicted to have increased binding affinity for hACE2.

| Amino acid change(s) | SARS-CoV-2 variant lineage<sup>78</sup> | Experimental $K_{D,app}$ ratio | NN_MM-GBSA $K_{D,app}$ ratio |
| --- | --- | --- | --- |
| K417T | P.1<sup>6</sup> | 0.55 | 0.15 |
| K417N | B.1.351<sup>5</sup> | 0.35 | 0.13 |
| L452R | B.1.429<sup>79</sup> | 1.05 | 1.19 |
| Y453F* | B.1.1.298<sup>80</sup> | 1.78 | 1.40 |
| S477N | B.1.526<sup>81#</sup> | 1.15 | 1.24 |
| E484K | B.1.351<sup>5</sup>, P.1<sup>6</sup>,<br>B.1.1.7<sup>4#</sup> | 1.15 | 1.32 |
| S494P | B.1.1.7<sup>4#</sup> | 1.00 | 1.03 |
| N501Y* | B.1.351<sup>5</sup>, B.1.1.7<sup>4</sup>,<br>P.1<sup>6</sup> | 1.74 | 1.37 |
| E484K+N501Y | P.1<sup>6</sup>, B.1.351<sup>5</sup> | — | 1.36 |
| E484K+S477N | B.1.526<sup>81#</sup> | — | 1.26 |
| E484K+N501Y+K417T | P.1<sup>6</sup> | — | 1.36 |
| E484K+N501Y+K417N | B.1.351<sup>5</sup> | — | 1.35 |
| V503W+E406F | De novo | — | 1.35 |
| V503W+Y505W* | De novo | — | 1.29 |

\* single amino-acid changes part of the training data

## Results

### Dataset preparation

The three-dimensional coordinates of the SARS-CoV-2 receptor binding domain in complex with human ACE2 were obtained from PDB entry 6LZG^65^. There exist 20 RBD residues contacting directly with hACE2 and making strong interactions at the binding interface.^33^ This gives a total of 380 possible single amino-acid variants by changing each one of the 20 RBD residues into the remaining 19 amino-acids. Of these, we chose all 27 variants with an increased binding affinity and 54 variants with lower binding affinity compared to the WT. The dataset was balanced by adding another 27 variants that exhibited binding enhancement though not in direct contact with hACE2 (see Supp. Table 1 for details). The selected 108 variants formed the training dataset in this study (see Figure 1). All RBD variants in the dataset were computationally modeled using Rosetta^48,49^ and analyzed for changes in binding affinity with hACE2 compared to the WT RBD. Experimental data on variant binding affinities were obtained from the deep mutagenesis study by Starr et. al.^33^ The study reported apparent dissociation constant *K*_D,app_ ratios for all possible variants with single-amino acid changes at every RBD position. A *K*_D,app_ ratio (i.e., *K*_D,app,WT_/*K*_D,app,variant_) for a variant greater than one implies stronger binding compared to WT, whereas a value less than one implies weaker binding (see also Figure 1C). *K*_D,app_ ratios can be related to changes in the free energy of binding (i.e., ΔΔ*G*_*bind*_) as (*K*_D,app*,variant*_)/(*K*_D,app*,WT*_) = exp(−ΔΔ*G*_*bind*_/RT). This enables direct comparison of experimental measurements with estimates in changes of free energy of binding from MM-GBSA and other computational methods (see Methods for details).

**Figure 1.**
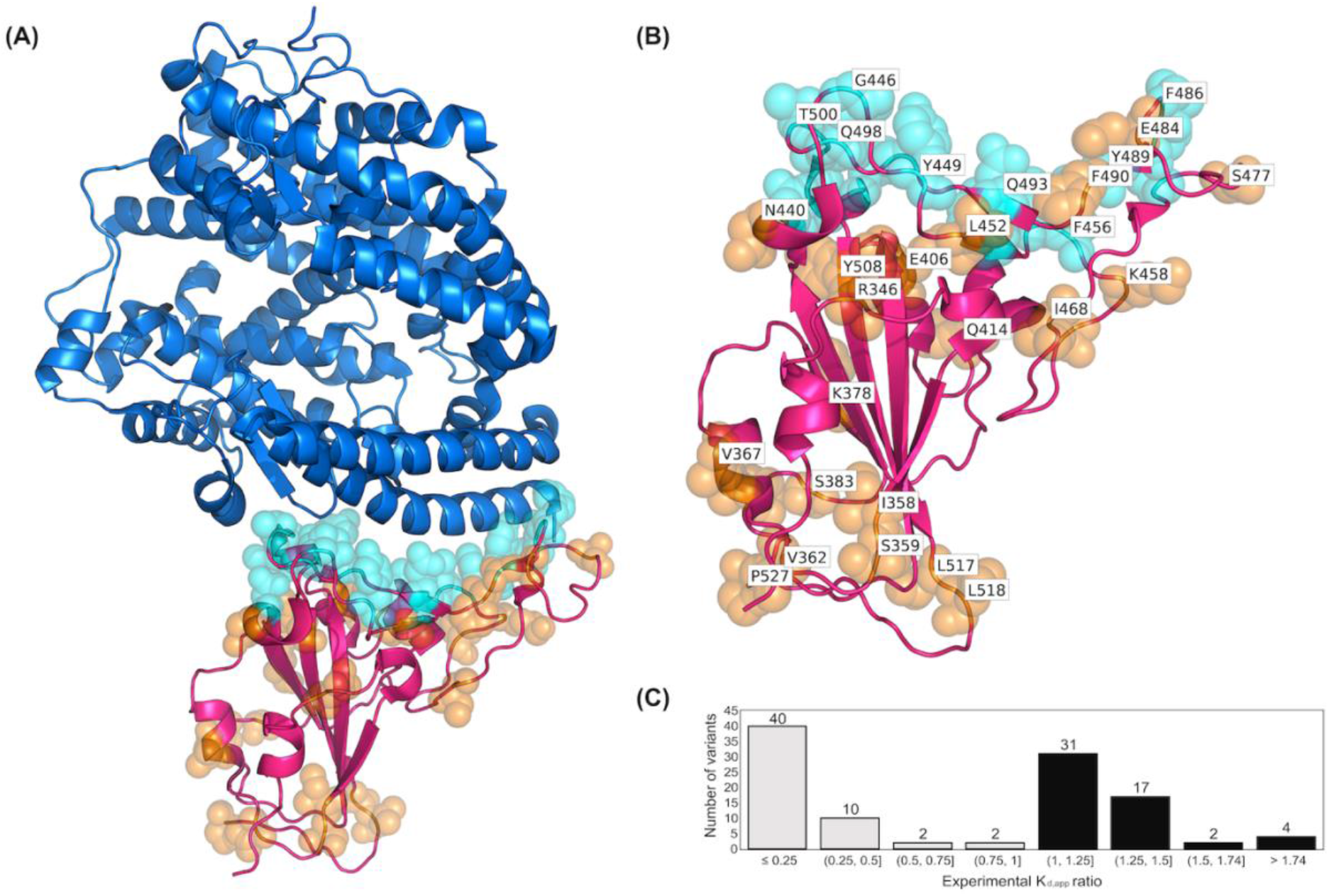
(A) The crystal structure of complex formed between RBD and hACE2 complex. The ACE2 protein is shown as a surface representation in blue and the RBD is shown in magenta. (B) Residues of the RBD variants that are in direct contact with hACE2 are depicted as cyan spheres. Residues that are not in direct contact are orange. (C) Histogram showing experimental *K*_*d,app*_ ratios for all 108 RBD variants in the dataset. The histogram bars in black denote number of variants in the training set with increasing binding affinity compared to WT (*K*_*d,app*_ ratio*>*1.0) and the bars in gray indicate the variant counts with decreasing binding affinity (*K*_*d,app*_ ratio *<* 1.0).

### Binding affinity change prediction for variants using MM-GBSA values from MD simulations

For each RBD variant, we first performed MD simulation of the hACE2-RBD complex followed by MM-GBSA analysis on frames derived from the simulation to calculate binding energies. For each variant, 48 independent initial configurations of the complex are generated by Monte Carlo minimization^66^ (see Methods). Starting from each configuration, a 4ns unconstrained MD simulation was carried out which generated a total of 192ns of MD trajectory for each variant. We used a sequence of short simulations, starting from several independent configurations instead of one long simulation trajectory - as it led to faster convergence, as also reported previously.^67^ 3D coordinates of the complex were extracted from each trajectory upon removing the solvent molecules after every 0.1ns, generating 1,920 different frames for each variant. MM-GBSA energies were calculated for all frames (see Methods) and subsequently averaged in 240 bins chosen randomly to obtain an ensemble of eight binding energy predictions for each variant. The median value of the ensemble of eight predictions was then chosen as the predicted binding energy Δ*G*_variant_ of the variant. The binding energy change for each variant ΔΔ*G*_*bind*_ is then obtained by subtracting the binding energy of the WT RBD-hACE2 complex Δ*G*_WT_. A negative ΔΔ*G*_*bind*_ value (corresponds to *K*_D,app_ ratio > 1) indicates improved binding affinity with hACE2, whereas a positive ΔΔ*G*_*bind*_ value (corresponding to *K*_D,app_ ratio < 1) implies lower binding affinity.

Using this computational workflow, we calculated the ΔΔ*G*_*bind*_ for the balanced dataset of 108 RBD variants. We score classification predictions using the percent recovery of correct variant classification (%VC) in terms of the direction of change in the binding affinity compared to WT. Note that a balanced training set was maintained, so as to alleviate the risk of biased predictions due to having more variants with worsening or improving binding affinities. We score quantitative binding affinity prediction using the Pearson correlation coefficient *r* (see Methods) between predicted and experimental ΔΔ*G*_*bind*_ values. We find that (see also Table 1) Rosetta slightly outperforms MM-GBSA in both prediction of the direction of change (i.e., %VC) and r value. This could be because of the scaling of the energy function within MM-GBSA not being in line with values obtained from experimental measurements leading to some outliers having very large predicted values (see Figure 2), causing a lower r value (i.e., r=0.33) than Rosetta (i.e., 0.47). In addition to the energy function from Rosetta^49^, three other computational servers were tested for the prediction; mCSM-PPI2^68^ utilizing graph-based signatures, the random forest model MutaBind2^69^ trained with Molecular Mechanics energies^70^ and evolutionary scores^71^, and SAAMBE-3d^72^ which uses a machine learning model trained on structural features. Using MutaBind2 and mCSM-PPI2, the performance in both %VC and r-value was worse than that of MM-GBSA and Rosetta. The predictions from SAAMBE-3d had a good correlation value r but were very poor in %VC (=0.53), almost the same as random prediction. This may be due to the fact that Rosetta and MM-GBSA attain a higher fidelity in the description of the underlying biophysics by using a detailed fully atomistic description of interactions and hence are better at distinguishing improving vs. worsening variants. Note that, since the numerical values ΔΔ*G*_*bind*_ for variants improving the binding affinity are quite small (maximum of ~ −0.3 kcal/mol) compared to those worsening the binding affinity (maximum of ~ +2.5kcal/mol), both metrics need to be simultaneously high to indicate robust prediction. Nevertheless, prediction metrics %VC and r calculated for MM-GBSA (or Rosetta) do not attain values that reflect reliable quantitative prediction. We next focused on improving prediction fidelity. This was accomplished by not merely using various energy terms in an additive fashion to assemble the overall binding energy, but instead by relying on a neural network to construct a nonlinear reassortment of these energy terms.

**Figure 2.**
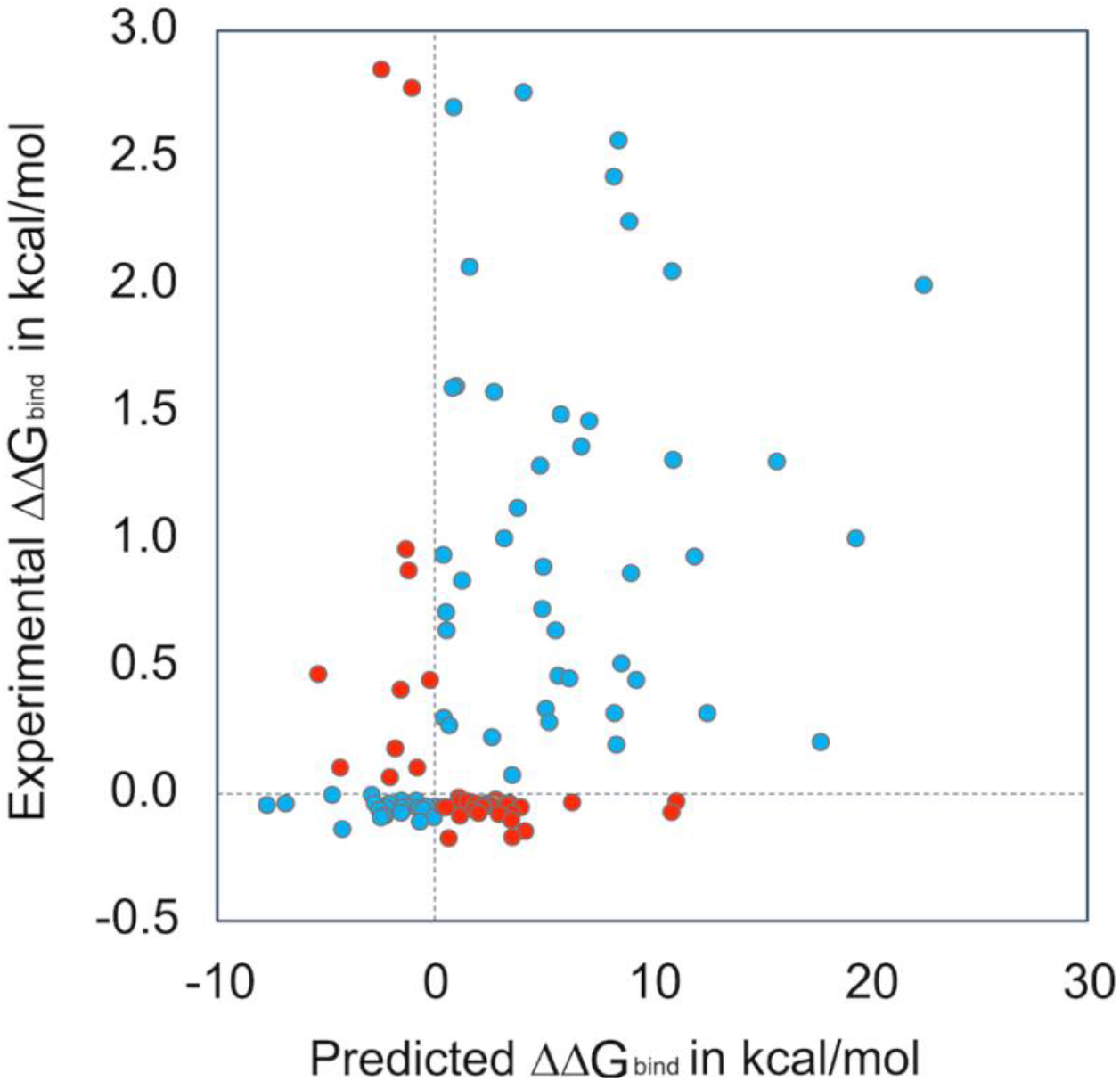
ΔΔ*G*_*bind*_ prediction performance of MM-GBSA binding energies on 108 RBD variants. Dotted horizontal and vertical lines are drawn for reference at experimental and predicted ΔΔ*G*_*bind*_ = 0. Shown in blue are variants for which the effect on binding affinity (sign of ΔΔ*G*_*bind*_) is predicted correctly compared to the experimental value. Those predicted incorrectly are shown in red.

### Neural network regression model trained on MM-GBSA energies and experimental K_d,app_ ratios

A neural network (NN) regression model with a single input and single output layer was built, targeting quantitative prediction of the *K*_*d,app*_ values for the 108 RBD variants. The NN had four fully connected hidden layers with 144 nodes in each layer (see Methods for details). The MM-GBSA ensemble of energies obtained from the MD trajectories of each variant was fed as input features. Each variant entry in the training set contains all 18 MM-GBSA energy terms (see Methods for description of terms) replicated for all eight sets of the ensemble. After passing through the hidden and output layers, each of the eight sets of energies outputs a single predicted value for *K*_*d,app*_. The model was trained to minimize the mean sum of squared error between the predicted *K*_*d,app*_ and experimental *K*_*d,app*_ values (see Methods for details). When making a prediction, each of the eight sets of energies is fed into the trained model to get a single *K*_*d,app*_ prediction and the final prediction value is recovered by taking the median of predictions from all eight ensembles. The overall computational workflow is summarized in Figure 3.

**Figure 3.**
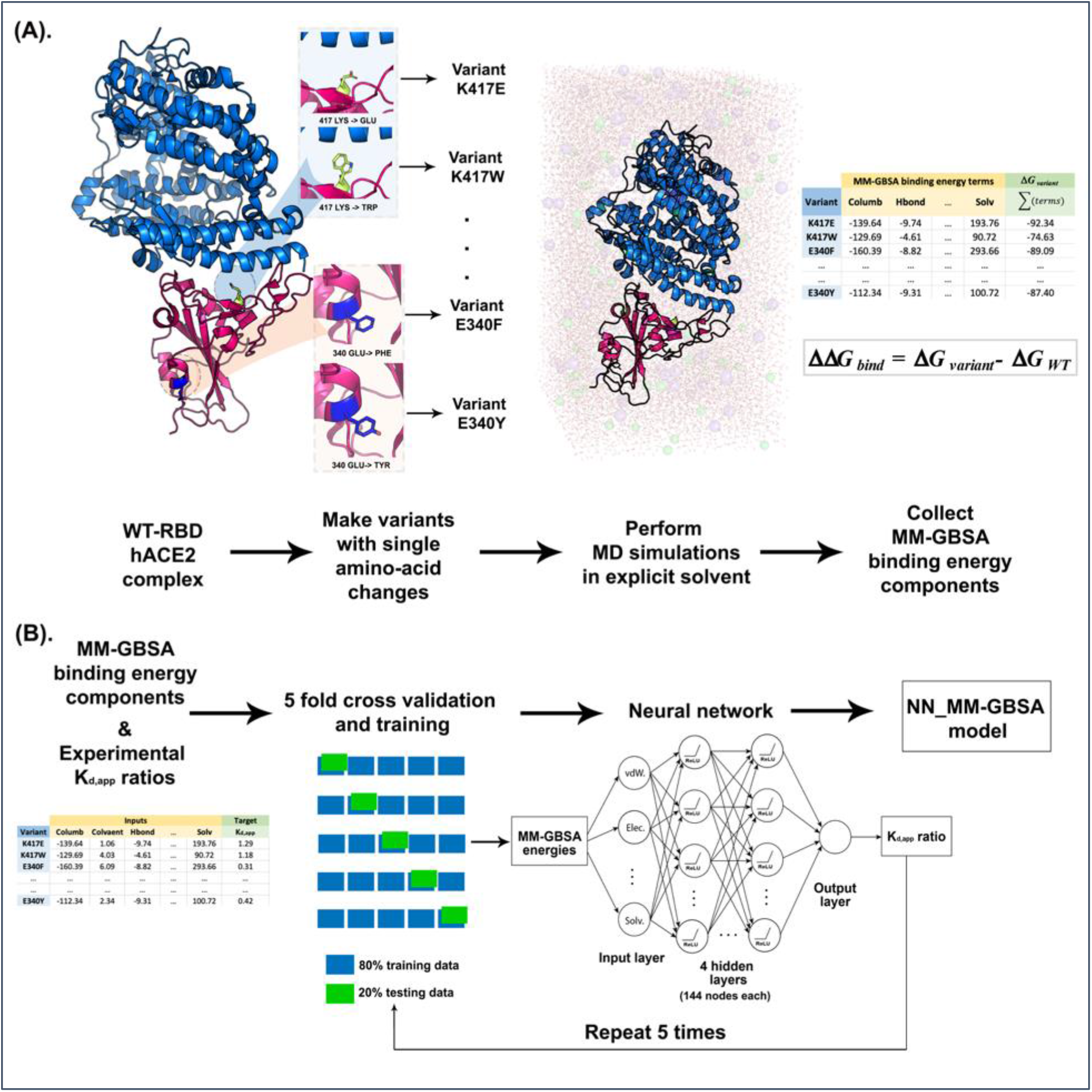
Schematic representation of the workflow for building NN_MM-GBSA model. (A) MD simulations were performed for each single point amino acid substitution variant in explicit solvent followed by MM-GBSA analysis to calculate the decomposed components of binding energies. (B) MM-GBSA binding energy components were fed as inputs to the Neural network with the experimental K_d,app_ ratios as the regression target. The model is trained using five cycles of the five-fold cross validation procedure.

During training and assessment of the model, a five-fold cross validation procedure was followed. During each cross-validation cycle, the 108 variants are randomly assigned to five groupings of approximately equal size. Four of these subsets are used as the training sets whereas the fifth becomes the testing set. This approach was chosen so that the testing set used to assess the prediction performance of the NN model is never used to train the predicting NN model. This 5-fold cross-validation was repeated 5 times using random assignments for the testing set. This led to the construction of 5×5 = 25 independently trained NN models with an average value of r=0.69 (obtained across the 25 models) and a standard deviation of only 0.03 implying both robust and accurate prediction (see Figure 4). Notably, the correlation coefficient of prediction improved by more than 2-fold compared to the MM-GBSA method (r=0.33) indicating that a higher order nonlinear structure, relevant to ΔΔ*G*_*bind*_ prediction embedded in the energy terms, was captured by the NN model. Correct variant classification (i.e., %VC) was also improved from 64% to 82% (see also Table 1).

**Figure 4.**
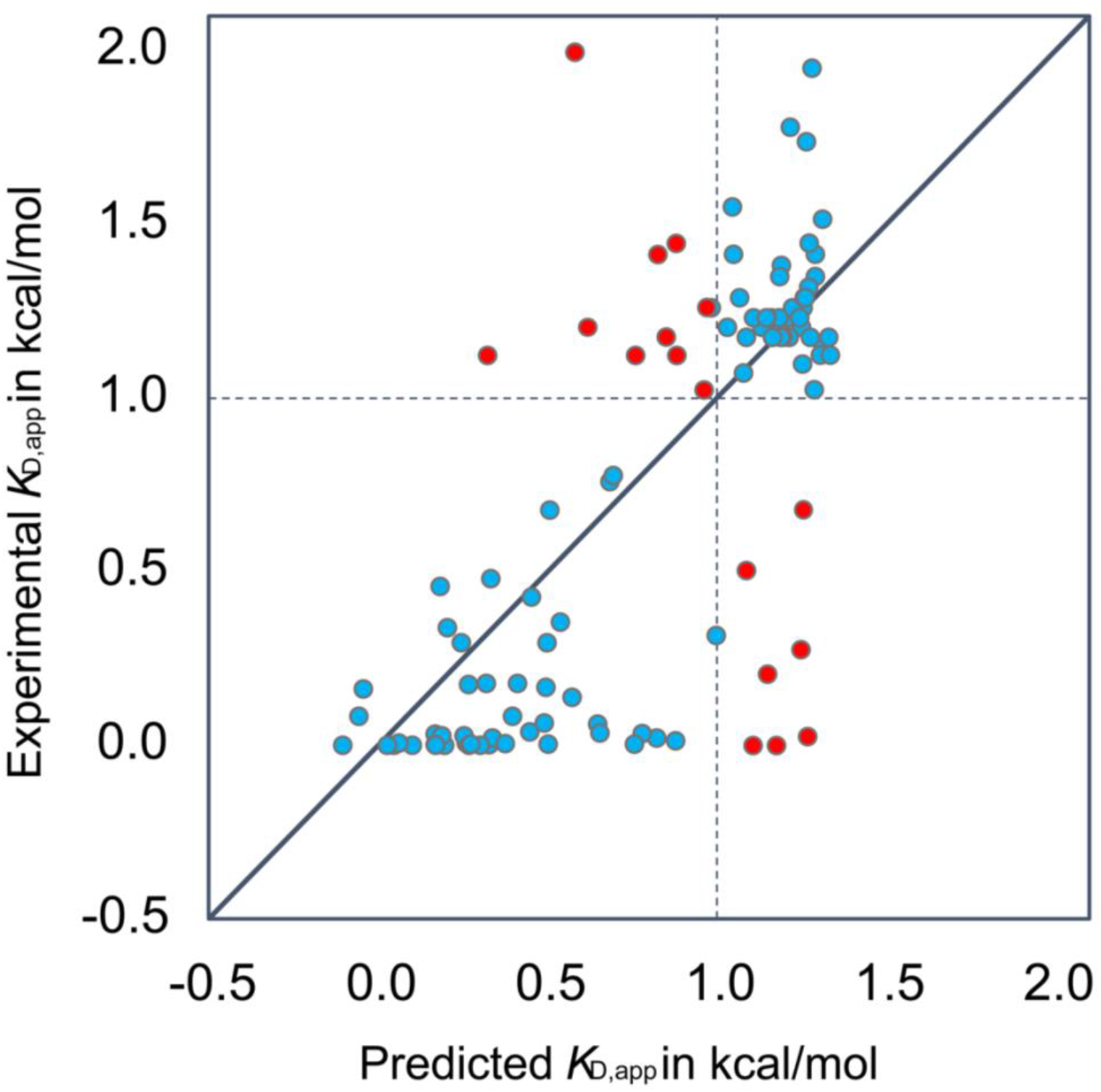
*K*_D,app_ ratios predicted by NN_MM-GBSA vs. experimental ratios obtained using five cycles of five-fold cross-validation study on the 108 RBD variants. An average correlation coefficient of r=0.69 (std. dev.=0.03) and average %VC of 82% (std. dev. = 0.02) were achieved. The Mean Squared Error was 0.20 (std. dev. = 0.02). The solid diagonal line y=x and the dotted horizontal and vertical lines at experimental and predicted *K*_D,app_ ratio = 1 are drawn for reference. Shown in blue are variants that were correctly classified as improving (or worsening) binding affinity with hACE2 and in red are the ones that were misclassified.

As a methodological check, we also explored whether the nonlinear nature of the NN model is needed to reach the gains in prediction or whether a linear regression model could simply re-weight the energy terms in a linear fashion and achieve similar performance. We found that a linear regression model improved the correlation coefficient r from 0.33 to 0.53 in comparison with the MM-GBSA prediction method but lowered %VC from 61% to 59%. This implies that the higher-order nonlinear re-assortment of energy terms is required for reaching improved prediction fidelity. As a follow up, we also explored if the energy terms from Rosetta^49,59^ could be used instead of the ones from MM-GBSA to construct a NN model of equivalent predictive ability. Note that the Rosetta energy function captures solvation effects implicitly without an explicit treatment of water molecules. We found that the gains in r and %VC for a NN model trained on the Rosetta energy terms were less than those seen when trained on MM-GBSA energies (i.e., r = 0.57, %VC=74.63). Both these analyses suggest that (1) explicit water treatment in MD simulations is important for correctly describing water-mediated hydrogen bonding and other electrostatic contacts at the interface while also enabling the sampling of a larger conformational landscape, which is necessary for capturing binding affinity changes due to nonlocal structural changes,^73^ and (2) the higher-order nonlinear regression afforded by a NN_MM-GBSA model are required to reach high prediction performance.

As a further demonstration that the NN_MM-GBSA model captures variant-specific information and does not simply carry-out numerical fitting, we performed a data scrambling test. Specifically, we re-assigned the variant definition (i.e., corresponding amino-acid change) to randomly chosen input energy terms, thereby destroying any variant-specific correspondence with the input features. We gradually increased the fraction of data scrambled and re-evaluated NN_MM-GBSA model performance. We found that as the fraction of scrambled data increases, the performance of the NN_MM-GBSA model declines. The %VC drops from the original 82.24% to 49.74% (almost completely random). This test reaffirms that the NN_MM-GBSA model indeed captured variant-specific information.

### NN_MM-GBSA model prediction of K_D,app_ ratios of circulating SARS-CoV-2 strains

Several amino-acid changes have been identified in the spike protein of several of the currently circulating SARS-CoV-2 variants.^74^ Some of the underlying single amino acid mutations were part of our balanced training set (i.e., N501Y, Y453F) whereas others (i.e., K417T, K417N, E484K, S477N, L452R,S494P) were not. The predicted *K*_D,app_ ratios along with experimental values (when available) and lineage names of variants that contain the corresponding amino acid changes are tabulated in Table 2. In all cases, amino acid changes were correctly classified as improving or worsening (i.e., %VC=100) with a good quantitative agreement (see Table 2). Notably, variant B.1.351 has the amino acid change N501Y first seen in B.1.1.7 along with additional changes E484K and K417N in the spike. The E484K change has been shown to be responsible for evasion of neutralization by several antibodies^75,76^ whereas the N501Y change has been associated with increased binding affinity to hACE2^33^ and increased transmission.^77^ Both amino acid changes were correctly classified by NN_MM-GBSA as improving binding affinity (see Table 2). In Table 2, we also include predictions for multiple simultaneous amino acid changes present in some circulating variants, alas experimental values are not available to this date which prevents any direct comparison. Nevertheless, significantly higher binding affinities were predicted for all tested double and triple amino acid variants present in currently circulating isolates (i.e., *K*_D,app_ ratios greater than 1.26 in all four cases). In addition, we included novel predicted variants V503W+E406F and V503W+Y505W, which we arrived at by maximizing binding affinity based on an exhaustive evaluation using the less accurate but more computationally tractable Rosetta energy function.^49^ Upon re-scoring using NN_MM-GBSA, we recovered high *K*_D,app_ ratios (i.e., 1.28 and 1.23, see Table 2). It is also worth noting that the single amino acid changes E406F and V503W by themselves cause a decline in binding affinity (experimental *K*_D,app_ ratio of 0.13 and 0.81) but when applied in combination lead to binding energy gains. This is probably due to a cooperative hydrophobic effect formed at the binding interface.

## Discussion

NN_MM-GBSA is a two-step procedure that uses the energy terms calculated for SARS-CoV-2 RBD variants from MM-GBSA to train a neural network model for predicting the qualitative and quantitative effects of amino acid changes in the RBD of the spike protein on the binding affinity with hACE2. Using a balanced training set of 108 variants, the method achieves a Pearson correlation coefficient of 0.69 between predicted and experimental values for the *K*_D,app_ ratios. In addition, the recovery of the correct effect of an amino acid change (i.e., improving or worsening binding) was 82%. We find that prediction is quite robust in terms of selection of training/testing sets with a standard deviation for the prediction or r of only 0.03. Notably, Starr et. al^33^ exhaustively assessed a total of ~4,000 variants for their binding affinity whereas, in this study, we used only a small fraction of the dataset (108 variants). As we continued to add additional members to the training dataset of 108 variants, no clear improving (or worsening) trendline was observed. Clearly, prediction fidelity is better for amino acid changes in positions that include at least one variant in the balanced training dataset. We plan to continue expanding upon the list; however, most of the remaining variants involve single amino acid changes far away from the RBD-hACE2 binding interface and thus contribute progressively less to binding affinity. The true value of NN_MM-GBSA is not the assessment of variants with single amino acid changes, but the surveillance for multiple amino acid change variants. We predicted the change in binding affinity upon the amino acid changes E484K+N501Y+K417N present in B1.351 and E484K+N501Y+K417T present in P.1. We found that N_MM-GBSA predicts for both, a significantly increased affinity for hACE2 (see Table 2) suggesting that the effect of amino-acid changes E484K and N501Y dominate the effect of K417N or K417T which are both known to decrease the binding affinity by themselves (see Table2).

In addition, we explored the possibility of double amino acid variants with even higher gains in binding affinity. Two such candidates (V503W+E406F and V503W+Y505W) were identified through exhaustive enumeration, using the Rosetta energy scoring function.^49^ For both of them, method NN_MM-GBSA confirmed high *K*_D,app_ ratios. These variants introduce large hydrophobic amino acids at the interface to enhance binding affinity with hACE2. The importance of the hydrophobic effect in protein-protein binding has been long recognized.^82^ Specifically, it has also been shown that the RBD-hACE2 interface is dominated by networks of hydrophobic contacts forming strong anchors for binding.^83^

A drawback of NN_MM-GBSA is that it requires *a priori* MD simulation of the variant under evaluation and collection of all energy terms using MM-GBSA analysis. This is computationally costly as a single calculation requires on average a total of 48 GPU-hours and 3 CPU-hours. Ideally, one could simply use existing energy terms generated from the balanced training set of 108 variants to make predictions for novel variants. However, this would require training a neural network model with more than just energy terms as descriptors. The use of sequence and/or structural features could provide a tractable path forward in this direction.

In principle, NN_MM-GBSA can also be used to assess infection potential for other non-human hosts by SARS-CoV-2 by assessing the binding energies of the spike RBD with the animal ACE2 receptors. The structures of non-human ACE2 are generally unavailable (except for bats^84^ and felines^85^) therefore, the first step would require homology modeling of the ACE2 receptor for the examined species. Efforts along this direction have been carried out for livestock and companion animals^32^ and assessments for high-risk animals are urgently needed. Assuming that training of NN_MM-GBSA using hACE2 data is robust, it could in principle be used to prospectively assess, the relative affinity of the RBD of circulating variants for various animal ACE2s. Crucially, this may provide the first indication that the viral variants have been established in non-human animal reservoir population(s). These variants can include both those already circulating and ones *de novo* predicted.

## Methods

### Rosetta calculations for independent structure generation

The 3D coordinates of SARS-CoV-2 viral spike RBD in complex with hACE2 were extracted from the crystal structure with PDB-id 6LZG.^65^ The obtained WT model was first pre-processed by removing all solvent molecules and all non-amino acid residues. For each of the 108 RBD variants with single point amino acid changes, 3D coordinates were generated using Rosettascripts^86^. First, the *PackRotamers* mover was used to build the variants with amino-acid changes and repack the rotamers. Then, for each variant, 48 independent configurations for MD simulations were generated using the *Relax*^66^ energy minimization protocol.

### Molecular dynamics simulations and MM-GBSA analysis

For each variant, the 3D coordinates of 48 independent configurations obtained using Rosetta (as described above) were prepared using protein preparation wizard^87^ protocol of Maestro in Schrödinger suite (v2019.4). Each configuration was then solvated with water using the tip3p^64^ model in an orthorhombic box with 10 Å buffer distance in each dimension. The residual charges were neutralized by adding Na^+^ and Cl^−^ ions at a salt concentration of 0.15 M. The solvated systems were minimized and pre-equilibrated using the default relaxation protocol of Desmond^88^ followed by a 4-ns production run using the amber99sb-ildn^63^ force field at 300 K and 1 atm. The NPT ensemble with periodic boundary conditions using particle mesh Ewald^89^ for long-range interactions. A time step of 2.0 fs was used and a cutoff distance of 9.0 Å was chosen for non-bonded interactions.

For each variant, the 4-ns trajectory for each of the 48 configurations was sampled at an interval of 0.1 ns, generating 1920 snapshots in total. For each snapshot, the Prime/MM-GBSA analysis^62^ was performed using *thermal_mmgbsa.py* script from the Schrödinger suite. The MM-GBSA analysis produces the binding energy and its constituent eight individual energy terms i.e., Coulomb energy, Covalent binding energy, van der Waals energy, Lipophilic energy, Generalized Born electrostatic solvation energy, Prime energy, Hydrogen-bonding correction, π-π packing correction, and Self-contact correction. Another set of values for these nine terms have also been calculated by not accounting for receptor and ligand conformational changes needed to form the complex. Due to a high degree of variation in the energies, we averaged data from 240 snapshots to produce a single set of averaged energy terms in the dataset. Thus, a total of 1920 snapshots generate eight sets of averaged energy terms for each variant. In total, these 18 energy values were utilized as the input features for NN construction.

### Neural network for MM-GBSA energies (NN_MM-GBSA)

A. Dataset Generation: MM-GBSA analysis was used to generate 18 energy components (as described above) which are fed as the input features for the NN_MM-GBSA model. Each input energy term across the whole data set was scaled independently to have zero mean and a variance of one. The output target was set to the experimental apparent dissociation constant *K*_D,app_ ratios, (*K*_D,app_)_variant_/(*K*_D,app_)_WT_. The experimental data for the 108 RBD variants (see Supp. Table 1 for list) was obtained from Starr et. al.^33^
B. Model Architecture: The neural network has a single input layer, a single output layer and four fully connected hidden layers with 144 nodes per layer, forming 18-(144-144-144-144)-1 structure. The rectified linear unit was used as the activation function for all of the hidden layers and the dropout regularization method was applied to hidden layers, with a dropout rate of 0.5 for the first and last hidden layers, and 0.75 for the rest.
C. Model Training: The model was trained through backpropagation to minimize the mean squared error between predicted K_D,app_ and target K_D,app_ values. Adam^90^ optimizer was used to perform the backpropagation with a learning rate of 0.001 and weight decay of 0.005. The training was performed for 2,000 epochs including the entire training data in each batch.
D. Model Evaluation: We used a five-fold cross validation protocol to evaluate the model, by splitting the whole dataset into 5 subsets. In one complete evaluation cycle, each of the 5 subsets was used as a test set once and the rest constituted the training set and a total of 5 such cycles were performed. Two metrics were employed to quantify the performance of NN_MM-GBSA model predictions: % correct variant classification (%VC), and the Pearson correlation coefficient (*r*). %VC is the percentage of instances in which a variant is classified correctly as increasing or decreasing the binding affinity compared to WT. The Pearson’s correlation coefficient is defined as:

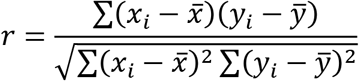

where *x*_*i*_ and *y*_*i*_ are the target and prediction for the *i*^th^ sample, and 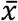 and 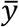 are the mean value for all *x*_*i*_ targets and *y*_*i*_ predictions.
E. Model Predictions: The NN_MM-GBSA model predictions are based on a single model trained using 100% of the training data. When making the prediction for a variant, the ensemble of eight sets of MM-GBSA energies are collected and each set is used to make a single prediction for *K*_D,app_ using the model. The median of eight predictions is the final predictor of the *K*_D,app_ of the variant.
F. Implementation: All codes were developed in Python using the PyTorch library.

### Rosetta calculations for ΔΔG_bind_ prediction

The complexes for 108 RBD variants were subject to *Relax*^66^ with harmonic constraints to prevent the structure from deviating significantly from the crystal structure. During *Relax*, rotamers of amino acid residues within 8 Å of the mutated amino acid were only allowed to repack (local packing). All default parameters were used for *Relax* with the ref2015 energy function.^49^. At the end of *Relax,* a gradient minimization is performed using *lbfgs*_*armijo* algorithm for 2000 steps after which the relevant metrics of binding were calculated using *InterfaceAnalyzer.*^91^ The binding energy, Δ*G*_*variant*_ of each variant is calculated as the average of *dG_separated* scores obtained from 30 independent *Relax* simulations. For each variant, a WT binding energy, Δ*G*_*WT*_ is calculated using the same protocol by making a dummy amino acid change (change amino acid to itself). Finally, the change in binding energy ΔΔ*G*_*bind*_ is calculated as ΔΔ*G*_*bind*_ = Δ*G*_*variant*_ − Δ*G*_*WT*_.

## Author contributions

CC, VSB, and CDM conceived, and designed the study. CC performed the MD simulations and built the NN models. VSB performed Rosetta calculations. DB built the linear regression models. All the authors wrote and approved the study.

## Acknowledgements

This activity was primarily supported by the United States Department of Agriculture (USDA) NIFA Award 2020-67015-32175 and also partially enabled by funding provided by The Center for Bioenergy Innovation a U.S. Department of Energy Research Center supported by the Office of Biological and Environmental Research in the DOE Office of Science (DE-AC05-000R22725).

Computations for this research were performed on the Pennsylvania State University’s Institute for Computational and Data Sciences’ Roar supercomputer. We also acknowledge Seed grant funding from the Penn State Huck Institutes of life sciences (to SVK).

Kurt Vandegrift was partially supported by a National Science Foundation Ecology and Evolution of Infectious Diseases program grant (# 1619072).

## Author Approvals

All the authors approve of the work.

## Appendix A. Supplementary data

Supplementary data to this article are given in supp_file.pdf

## References

1. McKee, M. & Stuckler, D. If the world fails to protect the economy, COVID-19 will damage health not just now but also in the future. Nature Medicine vol. 26 640–642 (2020).

2. Dobso, A. P. et al. Ecology and economics for pandemic prevention: Investments to prevent tropical deforestation and to limit wildlife trade will protect against future zoonosis outbreaks. Science vol. 369 379–381 (2020).

3. Horby, A. P. et al. Presented to SAGE on 21/1/21 Authors - Peter Horby, Catherine Huntley, Nick Davies, John Edmunds, Neil Ferguson, Graham Medley, Calum Semple.

4. Rambaut, A. et al. Preliminary genomic characterisation of an emergent SARS-CoV-2 lineage in the UK defined by a novel set of spike mutations - SARS-CoV-2 coronavirus / nCoV-2019 Genomic Epidemiology - Virological. Virological.org 1–9 (2020).

5. Tegally, H. et al. Emergence and rapid spread of a new severe acute respiratory syndrome-related coronavirus 2 (SARS-CoV-2) lineage with multiple spike mutations in South Africa. medRxiv (2020) doi:10.1101/2020.12.21.20248640.

6. Faria, N. R. et al. Genomic characterisation of an emergent SARS-CoV-2 lineage in Manaus: preliminary findings. Virological.Org 1–9 (2021).

7. Callaway, E. Fast-spreading COVID variant can elude immune responses. Nature 589, 500–501 (2021).

8. Ozono, S. et al. SARS-CoV-2 D614G spike mutation increases entry efficiency with enhanced ACE2-binding affinity. Nat. Commun. 12, 1–9 (2021).

9. Weissman, D. et al. D614G Spike Mutation Increases SARS CoV-2 Susceptibility to Neutralization. Cell Host Microbe 29, 23–31.e4 (2021).

10. Liu, Y. et al. Neutralizing Activity of BNT162b2-Elicited Serum - Preliminary Report. N. Engl. J. Med. (2021) doi:10.1056/NEJMc2102017.

11. Wang, Z. et al. mRNA vaccine-elicited antibodies to SARS-CoV-2 and circulating variants 2 3 4. bioRxiv 2021.01.15.426911 (2021) doi:10.1101/2021.01.15.426911.

12. Wu, K. et al. mRNA-1273 vaccine induces neutralizing antibodies against spike mutants from global SARS-CoV-2 variants. bioRxiv Prepr. Serv. Biol. 2021.01.25.427948 (2021) doi:10.1101/2021.01.25.427948.

13. Kim, D. et al. The Architecture of SARS-CoV-2 Transcriptome. Cell 181, 914–921.e10 (2020).

14. Cann, A. J. Principles of Molecular Virology: Sixth Edition. Principles of Molecular Virology: Sixth Edition (2015). doi:10.1016/C2014-0-01081-7.

15. Mercatelli, D. & Giorgi, F. M. Geographic and Genomic Distribution of SARS-CoV-2 Mutations. Front. Microbiol. 11, 1800 (2020).

16. Callaway, E. The coronavirus is mutating - does it matter? Nature vol. 585 174–177 (2020).

17. Abdelrahman, Z., Li, M. & Wang, X. Comparative Review of SARS-CoV-2, SARS-CoV, MERS-CoV, and Influenza A Respiratory Viruses. Frontiers in Immunology vol. 11 552909 (2020).

18. Kim, Y. Il et al. Infection and Rapid Transmission of SARS-CoV-2 in Ferrets. Cell Host Microbe 27, 704–709.e2 (2020).

19. Koopmans, M. SARS-CoV-2 and the human-animal interface: outbreaks on mink farms. The Lancet Infectious Diseases vol. 21 18–19 (2021).

20. Montagutelli, X. et al. The B1.351 and P.1 variants extend SARS-CoV-2 host range to mice. bioRxiv 2021.03.18.436013 (2021) doi:10.1101/2021.03.18.436013.

21. Davies, N. G. et al. Estimated transmissibility and impact of SARS-CoV-2 lineage B.1.1.7 in England. Science (2021) doi:10.1126/science.abg3055.

22. Li, Q. et al. The Impact of Mutations in SARS-CoV-2 Spike on Viral Infectivity and Antigenicity. Cell 182, 1284–1294.e9 (2020).

23. Zhang, L. et al. SARS-CoV-2 spike-protein D614G mutation increases virion spike density and infectivity. Nat. Commun. 11, 1–9 (2020).

24. Zhou, W. et al. N439K variant in spike protein may alter the infection efficiency and antigenicity of SARS-CoV-2 based on molecular dynamics simulation. bioRxiv (2020) doi:10.1101/2020.11.21.392407.

25. Zhou, P. et al. A pneumonia outbreak associated with a new coronavirus of probable bat origin. Nature 579, 270–273 (2020).

26. Walls, A. C. et al. Structure, Function, and Antigenicity of the SARS-CoV-2 Spike Glycoprotein. Cell (2020) doi:10.1016/j.cell.2020.02.058.

27. Wan, Y., Shang, J., Graham, R., Baric, R. S. & Li, F. Receptor recognition by novel coronavirus from Wuhan: An analysis based on decade-long structural studies of SARS. J. Virol. (2020) doi:10.1128/jvi.00127-20.

28. Hwang, S. S. et al. MRNA destabilization by BTG1 and BTG2 maintains T cell quiescence. Science (80-.). 367, 1255–1260 (2020).

29. Letko, M., Marzi, A. & Munster, V. Functional assessment of cell entry and receptor usage for SARS-CoV-2 and other lineage B betacoronaviruses. Nat. Microbiol. 5, 562–569 (2020).

30. Li, W. et al. Angiotensin-converting enzyme 2 is a functional receptor for the SARS coronavirus. Nature 426, 450–454 (2003).

31. Hoffmann, M. et al. SARS-CoV-2 Cell Entry Depends on ACE2 and TMPRSS2 and Is Blocked by a Clinically Proven Protease Inhibitor. Cell 181, 271–280.e8 (2020).

32. Chowdhury, R., Boorla, V. S. & Maranas, C. D. Computational biophysical characterization of the SARS-CoV-2 spike protein binding with the ACE2 receptor and implications for infectivity. Comput. Struct. Biotechnol. J. 18, 2573–2582 (2020).

33. Starr, T. N. et al. Deep Mutational Scanning of SARS-CoV-2 Receptor Binding Domain Reveals Constraints on Folding and ACE2 Binding. Cell 182, 1295–1310.e20 (2020).

34. Guruprasad, L. Human SARS CoV-2 spike protein mutations. Proteins Struct. Funct. Bioinforma. (2021) doi:10.1002/prot.26042.

35. Piccoli, L. et al. Mapping Neutralizing and Immunodominant Sites on the SARS-CoV-2 Spike Receptor-Binding Domain by Structure-Guided High-Resolution Serology. Cell 183, 1024–1042.e21 (2020).

36. Starr, T. N. et al. Prospective mapping of viral mutations that escape antibodies used to treat COVID-19. Science 371, 850–854 (2021).

37. Greaney, A. J. et al. Complete Mapping of Mutations to the SARS-CoV-2 Spike Receptor-Binding Domain that Escape Antibody Recognition. Cell Host Microbe 29, 44–57.e9 (2021).

38. Bayarri-Olmos, R. et al. The SARS-CoV-2 Y453F mink variant displays a pronounced increase in ACE-2 affinity but does not challenge antibody neutralization. J. Biol. Chem. 0, 100536 (2021).

39. Baum, A. et al. REGN-COV2 antibodies prevent and treat SARS-CoV-2 infection in rhesus macaques and hamsters. Science (80-.). 370, 1110–1115 (2020).

40. Tang, T., Bidon, M., Jaimes, J. A., Whittaker, G. R. & Daniel, S. Coronavirus membrane fusion mechanism offers a potential target for antiviral development. Antiviral Research vol. 178 (2020).

41. Fehr, A. R. & Perlman, S. Coronaviruses: An overview of their replication and pathogenesis. Coronaviruses Methods Protoc. 1–23 (2015) doi:10.1007/978-1-4939-2438-7_1.

42. Mohammad, A. et al. Higher binding affinity of furin for SARS-CoV-2 spike (S) protein D614G mutant could be associated with higher SARS-CoV-2 infectivity. International Journal of Infectious Diseases vol. 103 611–616 (2021).

43. Weissman, D. et al. D614G Spike Mutation Increases SARS CoV-2 Susceptibility to Neutralization. Cell Host Microbe 29, 23–31.e4 (2021).

44. Gobeil, S. M. C. et al. D614G Mutation Alters SARS-CoV-2 Spike Conformation and Enhances Protease Cleavage at the S1/S2 Junction. Cell Rep. 34, (2021).

45. Chen, J., Wang, R., Wang, M. & Wei, G. W. Mutations Strengthened SARS-CoV-2 Infectivity. J. Mol. Biol. 432, 5212–5226 (2020).

46. Jankauskaite, J., Jiménez-García, B., Dapkunas, J., Fernández-Recio, J. & Moal, I. H. SKEMPI 2.0: An updated benchmark of changes in protein-protein binding energy, kinetics and thermodynamics upon mutation. Bioinformatics 35, 462–469 (2019).

47. Guerois, R., Nielsen, J. E. & Serrano, L. Predicting changes in the stability of proteins and protein complexes: A study of more than 1000 mutations. J. Mol. Biol. 320, 369–387 (2002).

48. Leaver-Fay, A. et al. Rosetta3: An object-oriented software suite for the simulation and design of macromolecules. in Methods in Enzymology vol. 487 545–574 (Academic Press Inc., 2011).

49. Alford, R. F. et al. The Rosetta All-Atom Energy Function for Macromolecular Modeling and Design. J. Chem. Theory Comput. 13, 3031–3048 (2017).

50. Beard, H., Cholleti, A., Pearlman, D., Sherman, W. & Loving, K. A. Applying Physics-Based Scoring to Calculate Free Energies of Binding for Single Amino Acid Mutations in Protein-Protein Complexes. PLoS One 8, e82849 (2013).

51. Petukh, M., Dai, L. & Alexov, E. SAAMBE: Webserver to predict the charge of binding free energy caused by amino acids mutations. Int. J. Mol. Sci. 17, (2016).

52. Xiong, P., Zhang, C., Zheng, W. & Zhang, Y. BindProfX: Assessing Mutation-Induced Binding Affinity Change by Protein Interface Profiles with Pseudo-Counts. J. Mol. Biol. 429, 426–434 (2017).

53. Dehouck, Y., Kwasigroch, J. M., Rooman, M. & Gilis, D. BeAtMuSiC: Prediction of changes in protein-protein binding affinity on mutations. Nucleic Acids Res. 41, (2013).

54. Moal, I. H. & Fernandez-Recio, J. Intermolecular contact potentials for protein-protein interactions extracted from binding free energy changes upon mutation. J. Chem. Theory Comput. 9, 3715–3727 (2013).

55. Barlow, K. A. et al. Flex ddG: Rosetta Ensemble-Based Estimation of Changes in Protein-Protein Binding Affinity upon Mutation. J. Phys. Chem. B 122, 5389–5399 (2018).

56. Shringari, S. R., Giannakoulias, S., Ferrie, J. J. & James Petersson, E. Rosetta custom score functions accurately predict ΔΔ: G of mutations at protein-protein interfaces using machine learning. Chem. Commun. 56, 6774–6777 (2020).

57. Wang, M., Cang, Z. & Wei, G.-W. A topology-based network tree for the prediction of protein–protein binding affinity changes following mutation. Nat. Mach. Intell. 2, 116–123 (2020).

58. Geng, C., Xue, L. C., Roel‐Touris, J. & Bonvin, A. M. J. J. Finding the ΔΔ *G* spot: Are predictors of binding affinity changes upon mutations in protein–protein interactions ready for it? WIREs Comput. Mol. Sci. 9, e1410 (2019).

59. Leaver-Fay, A. et al. Rosetta3: An object-oriented software suite for the simulation and design of macromolecules. in Methods in Enzymology vol. 487 545–574 (Academic Press Inc., 2011).

60. Jacobson, M. P. et al. A hierarchical approach to all-atom protein loop prediction. Proteins Struct. Funct. Bioinforma. 55, 351–367 (2004).

61. Jacobson, M. P., Friesner, R. A., Xiang, Z. & Honig, B. On the role of the crystal environment in determining protein side-chain conformations. J. Mol. Biol. 320, 597–608 (2002).

62. Li, J. et al. The VSGB 2.0 model: A next generation energy model for high resolution protein structure modeling. Proteins Struct. Funct. Bioinforma. 79, 2794–2812 (2011).

63. Lindorff-Larsen, K. et al. Improved side-chain torsion potentials for the Amber ff99SB protein force field. Proteins Struct. Funct. Bioinforma. 78, 1950–1958 (2010).

64. Jorgensen, W. L., Chandrasekhar, J., Madura, J. D., Impey, R. W. & Klein, M. L. Comparison of simple potential functions for simulating liquid water. J. Chem. Phys. 79, 926–935 (1983).

65. Wang, Q. et al. Structural and Functional Basis of SARS-CoV-2 Entry by Using Human ACE2. Cell 181, 894–904.e9 (2020).

66. Conway, P., Tyka, M. D., DiMaio, F., Konerding, D. E. & Baker, D. Relaxation of backbone bond geometry improves protein energy landscape modeling. Protein Sci. 23, 47–55 (2014).

67. Wang, E. et al. End-Point Binding Free Energy Calculation with MM/PBSA and MM/GBSA: Strategies and Applications in Drug Design. Chemical Reviews vol. 119 9478–9508 (2019).

68. Rodrigues, C. H. M., Myung, Y., Pires, D. E. V. & Ascher, D. B. MCSM-PPI2: predicting the effects of mutations on protein-protein interactions. Nucleic Acids Res. 47, W338–W344 (2019).

69. Zhang, N. et al. MutaBind2: Predicting the Impacts of Single and Multiple Mutations on Protein-Protein Interactions. iScience 23, 100939 (2020).

70. Huang, J. & Mackerell, A. D. CHARMM36 all-atom additive protein force field: Validation based on comparison to NMR data. J. Comput. Chem. 34, 2135–2145 (2013).

71. Choi, Y., Sims, G. E., Murphy, S., Miller, J. R. & Chan, A. P. Predicting the Functional Effect of Amino Acid Substitutions and Indels. PLoS One 7, e46688 (2012).

72. Pahari, S. et al. SAAMBE-3D: Predicting effect of mutations on protein–protein interactions. Int. J. Mol. Sci. 21, (2020).

73. Wang, E. et al. End-Point Binding Free Energy Calculation with MM/PBSA and MM/GBSA: Strategies and Applications in Drug Design. Chem. Rev. 119, 9478–9508 (2019).

74. Benton, D. J. et al. The effect of the D614G substitution on the structure of the spike glycoprotein of SARS-CoV-2. Proc. Natl. Acad. Sci. 118, e2022586118 (2021).

75. Baum, A. et al. Antibody cocktail to SARS-CoV-2 spike protein prevents rapid mutational escape seen with individual antibodies. Science (80-.). 369, 1014–1018 (2020).

76. Ku, Z. et al. Molecular determinants and mechanism for antibody cocktail preventing SARS-CoV-2 escape. Nat. Commun. 12, 1–13 (2021).

77. Zhao, S. et al. Quantifying the transmission advantage associated with N501Y substitution of SARS-CoV-2 in the UK: an early data-driven analysis. J. Travel Med. 28, 1–3 (2021).

78. Rambaut, A. et al. A dynamic nomenclature proposal for SARS-CoV-2 lineages to assist genomic epidemiology. Nat. Microbiol. 5, 1403–1407 (2020).

79. Zhang, W. et al. Emergence of a Novel SARS-CoV-2 Variant in Southern California. JAMA (2021) doi:10.1001/jama.2021.1612.

80. Oreshkova, N. et al. SARS-CoV-2 infection in farmed minks, the Netherlands, April and May 2020. Eurosurveillance 25, (2020).

81. West, A. P., Barnes, C. O., Yang, Z. & Bjorkman, P. J. SARS-CoV-2 lineage B.1.526 emerging in the New York region detected by software utility created to query the spike mutational landscape. bioRxiv 2021.02.14.431043 (2021) doi:10.1101/2021.02.14.431043.

82. Keskin, O., Gursoy, A., Ma, B. & Nussinov, R. Principles of protein-protein interactions: What are the preferred ways for proteins to interact? Chemical Reviews vol. 108 1225–1244 (2008).

83. Wang, Y., Liu, M. & Gao, J. Enhanced receptor binding of SARS-CoV-2 through networks of hydrogen-bonding and hydrophobic interactions. Proc. Natl. Acad. Sci. 117, 202008209 (2020).

84. Liu, K. et al. Cross-species recognition of SARS-CoV-2 to bat ACE2. Proc. Natl. Acad. Sci. U. S. A. 118, (2021).

85. Wu, L. et al. Broad host range of SARS-CoV-2 and the molecular basis for SARS-CoV-2 binding to cat ACE2. Cell Discov. 6, 1–12 (2020).

86. Fleishman, S. J. et al. RosettaScripts: A Scripting Language Interface to the Rosetta Macromolecular Modeling Suite. PLoS One 6, e20161 (2011).

87. Madhavi Sastry, G., Adzhigirey, M., Day, T., Annabhimoju, R. & Sherman, W. Protein and ligand preparation: Parameters, protocols, and influence on virtual screening enrichments. J. Comput. Aided. Mol. Des. 27, 221–234 (2013).

88. Bowers, K. J. et al. Scalable algorithms for molecular dynamics simulations on commodity clusters. Proc. 2006 ACM/IEEE Conf. Supercomput. SC’06 (2006) doi:10.1145/1188455.1188544.

89. Darden, T., York, D. & Pedersen, L. Particle mesh Ewald: An N·log(N) method for Ewald sums in large systems. J. Chem. Phys. 98, 10089–10092 (1993).

90. Kingma, D. P. & Ba, J. L. Adam: A method for stochastic optimization. in 3rd International Conference on Learning Representations, ICLR 2015 - Conference Track Proceedings (International Conference on Learning Representations, ICLR, 2015).

91. Benjamin Stranges, P. & Kuhlman, B. A comparison of successful and failed protein interface designs highlights the challenges of designing buried hydrogen bonds. Protein Sci. 22, 74–82 (2013).

