## Supplemental Table 1 for "Computational prediction of the effect of amino acid changes on the binding affinity between SARS-CoV-2 spike protein and the human ACE2 receptor"

**Supp. Table 1.** List of 108 RBD variants with single amino acid changes used in the study with their corresponding experimental values from Starr. et al.[1]

| Amino acid change | Experimental $K_{d,app}$ ratio |
|---|---|
| 498H | 2.00 |
| 501F | 1.95 |
| 453F | 1.78 |
| 501Y | 1.74 |
| 385R | 1.55 |
| 493M | 1.51 |
| 414A | 1.45 |
| 498Y | 1.45 |
| 484R | 1.41 |
| 498F | 1.41 |
| 501V | 1.41 |
| 367W | 1.38 |
| 493A | 1.35 |
| 505W | 1.35 |
| 517M | 1.35 |
| 493Y | 1.32 |
| 367A | 1.29 |
| 501W | 1.29 |
| 527M | 1.29 |
| 358F | 1.26 |
| 501T | 1.26 |
| 503K | 1.26 |
| 503M | 1.26 |
| 383E | 1.23 |
| 452K | 1.23 |
| 458D | 1.23 |
| 460K | 1.23 |
| 477D | 1.23 |
| 490K | 1.23 |
| 518S | 1.23 |
| 346H | 1.20 |
| 359Q | 1.20 |
| 362T | 1.20 |
| 366N | 1.20 |
| 378R | 1.20 |
| 453K | 1.20 |

| | |
|---|---|
| 367F | 1.17 |
| 406Q | 1.17 |
| 440K | 1.17 |
| 452Q | 1.17 |
| 468M | 1.17 |
| 498W | 1.17 |
| 508H | 1.17 |
| 527Q | 1.17 |
| 493F | 1.15 |
| 455M | 1.12 |
| 493K | 1.12 |
| 493L | 1.12 |
| 493V | 1.12 |
| 503I | 1.12 |
| 439K | 1.10 |
| 503L | 1.07 |
| 493G | 1.02 |
| 503R | 1.02 |
| 446N | 0.78 |
| 446E | 0.76 |
| 417R | 0.68 |
| 446Q | 0.68 |
| 453R | 0.50 |
| 417V | 0.48 |
| 486A | 0.46 |
| 453Q | 0.43 |
| 439Q | 0.35 |
| 455A | 0.34 |
| 439G | 0.32 |
| 486M | 0.30 |
| 496A | 0.30 |
| 500A | 0.28 |
| 456H | 0.20 |
| 417E | 0.18 |
| 439H | 0.18 |
| 455H | 0.17 |
| 455S | 0.17 |
| 501E | 0.16 |
| 475R | 0.14 |
| 449F | 0.08 |

| | |
|---|---|
| 487D | 0.08 |
| 500N | 0.06 |
| 449N | 0.06 |
| 449P | 0.04 |
| 496T | 0.03 |
| 498I | 0.03 |
| 455Y | 0.03 |
| 493D | 0.03 |
| 498L | 0.03 |
| 475P | 0.02 |
| 475Y | 0.02 |
| 486D | 0.02 |
| 500V | 0.01 |
| 456A | 0.01 |
| 487L | 0.01 |
| 496D | 0.01 |
| 487R | 0.01 |
| 505R | 0.00 |
| 505L | 0.00 |
| 439E | 0.00 |
| 456P | 0.00 |
| 502S | 0.00 |
| 489D | 0.00 |
| 505T | 0.00 |
| 453D | 0.00 |
| 489R | 0.00 |
| 505P | 0.00 |
| 489K | 0.00 |
| 502P | 0.00 |
| 505G | 0.00 |
| 502Q | 0.00 |
| 453P | 0.00 |

### References


1. Starr, T. N. *et al.* Deep Mutational Scanning of SARS-CoV-2 Receptor Binding Domain Reveals Constraints on Folding and ACE2 Binding. *Cell* **182**, 1295-1310.e20 (2020).
